# Visualization of Phase-Amplitude Coupling Using Rhythmic High-Frequency Activity

**DOI:** 10.1101/2020.08.12.247775

**Authors:** Hiroaki Hashimoto, Hui Ming Khoo, Takufumi Yanagisawa, Naoki Tani, Satoru Oshino, Haruhiko Kishima, Masayuki Hirata

**Affiliations:** Department of Neurological Diagnosis and Restoration, Graduate School of Medicine, Osaka University, Suita 565-0871, Japan; Department of Neurosurgery, Otemae Hospital, Osaka, 540-0008, Japan; Endowed Research Department of Clinical Neuroengineering, Global Center for Medical Engineering and Informatics, Osaka University, Suita 565-0871, Japan; Department of Neurosurgery, Graduate School of Medicine, Osaka University, Suita 565-0871, Japan

**Keywords:** video-recorded seizures, phase-amplitude coupling, theta band, high-frequency, intracranial EEG

## Abstract

**Objective:** High-frequency activities (HFAs) and phase-amplitude coupling (PAC) are gaining attention as key neurophysiological biomarkers for studying human epilepsy. We aimed to clarify and visualize how HFAs are modulated by the phase of low-frequency bands during seizures.

**Methods:** We used intracranial electrodes to record seizures of symptomatic focal epilepsy (15 seizures in seven patients). Ripples (80–250 Hz), as representative of HFAs, were evaluated along with PAC. The synchronization index (SI), representing PAC, was used to analyze the coupling between the amplitude of ripples and the phase of lower frequencies. We created a video in which the intracranial electrode contacts were represented by circles that were scaled linearly to the power changes of ripple.

**Results:** The main low frequency band modulating ictal-ripple activities was the θ band (4–8 Hz), and after completion of ictal-ripple burst, δ (1–4 Hz)-ripple PAC occurred. The video showed that fluctuation of the diameter of these circles indicated the rhythmic changes during significant high values of θ-ripple PAC.

**Conclusions:** We inferred that ripple activities occurring during seizure evolution were modulated by θ rhythm. In addition, we concluded that rhythmic circles’ fluctuation presented in the video represents the PAC phenomenon. Our video is thus a useful tool for understanding how ripple activity is modulated by the low-frequency phase in relation with PAC.

## Introduction

Intracranial electroencephalograms (iEEG) allow acquisition of wideband waveforms, from slow shift to high-frequency activities (HFAs), at a high signal-to-noise ratio. Direct current (DC) shifts and infraslow activities, which are very slow-frequency components, appear in the seizure onset zone (SOZ) during a seizure ^1–4^. HFAs can be physiological, i.e., those recorded during a task ^5^, or pathological, i.e., those observed during seizures or interictal period in epileptic patients ^2^ ^6^ ^7^. High frequency oscillations (HFOs) are subgroup of HFAs and isolated oscillations standing out from the background, within the high frequency range usually above 80 Hz. Ictal HFOs occur in the SOZ ^8^ ^9^, and HFOs can be further classified into ripples (80– 250 Hz) and fast ripples (250–500 Hz) ^9^. Previous study reported that HFAs were useful for detection of seizures^10^.

The amplitude of HFAs is modulated by the low-frequency oscillation phase ^11^. Physiologically, this phase-amplitude coupling (PAC) has various functional roles in cortical processing such as motor execution ^12^ and sensory processing ^13^. In the ictal state, PAC achieved high values in the SOZ ^14^ ^15^. Moreover, it is reported that PAC between the infraslow phase and HFAs amplitude preceded the seizure onset (SO) ^16^.

PAC achieves high values in the SOZ during seizures; however, there is no consensus about the main low frequency band that modulates the amplitude of HFAs. Several low frequency bands have been reported like δ ^15^ ^17^ ^18^, θ ^14^, α ^14^, and β ^19^ bands. Furthermore, whereas dynamic HFAs changes in the ictal state have been visualized using topographic videos ^6^, dynamic PAC changes have not been visualized. The purpose of this study is twofold. One is to clearly identify the main low frequency band contributing to PAC during seizures. The other is to visualize the PAC phenomenon. Using the iEEG, we collected data from 15 seizures in seven patients with medically refractory focal epilepsy. First, we evaluated dynamic changes and characteristics of PAC from pre-ictal to late-ictal. Next, we created a video for each patient in which circles, corresponding to electrodes, change their diameters that correlate linearly with the power of HFAs represented by ripples (Video S1–3). We hypothesized that PAC can be visualized using dynamic ripple power changes modulated by a low-frequency band.

## Methods

### Subjects

We included seven patients with drug-resistant focal epilepsy who underwent intracranial electrodes placement for presurgical invasive EEG study (one female, 15–47 years of age, mean age: 26 ± 12.1 years) (Table 1) who were admitted to Osaka University Hospital from July 2018 to July 2019. This retrospective study was approved by the Ethics Committee of Osaka University Hospital (Suita, Japan) (approval no., 19193). Informed consent was obtained by the opt-out method on our center’s website.

**Table 1.**
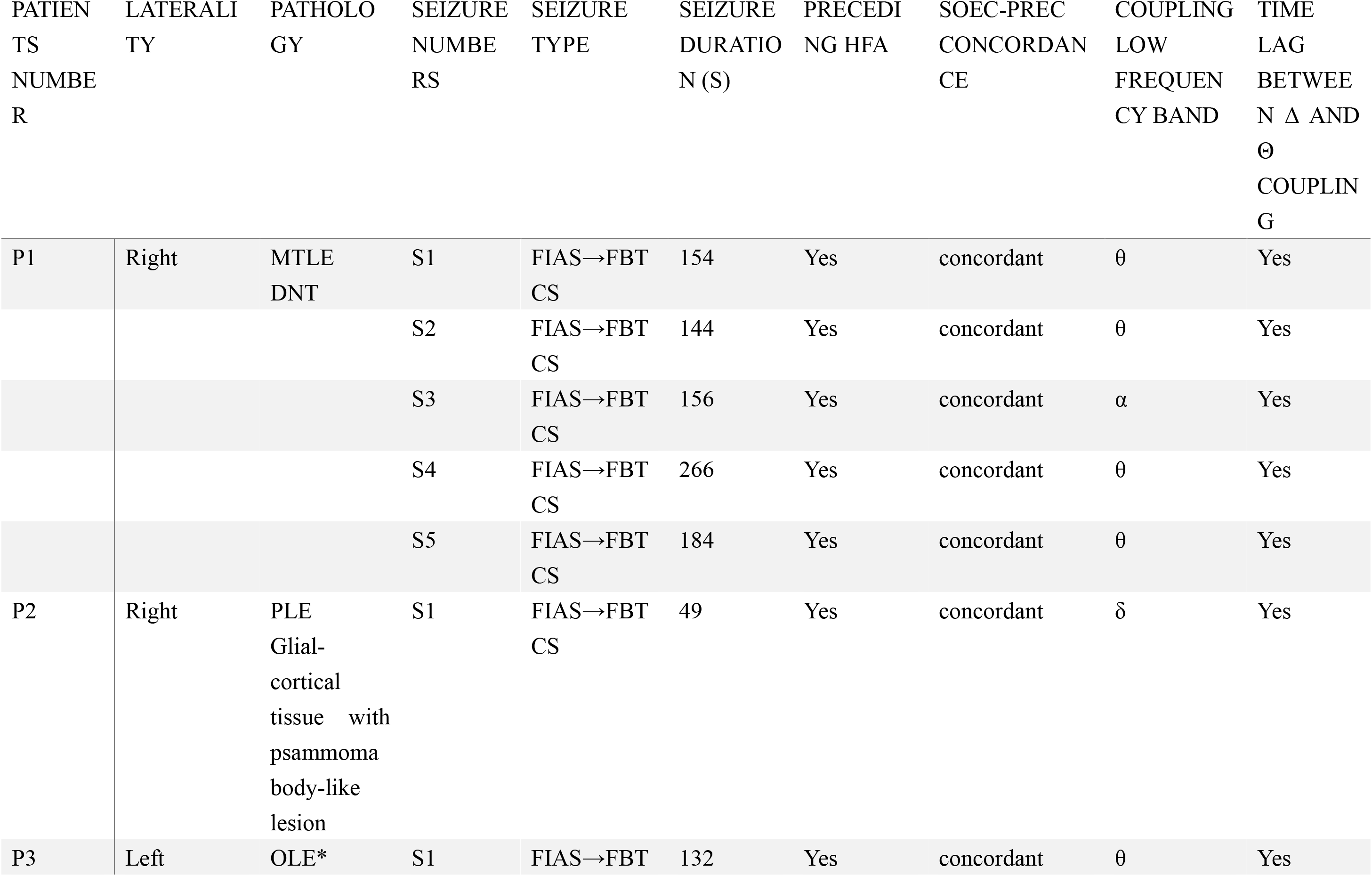
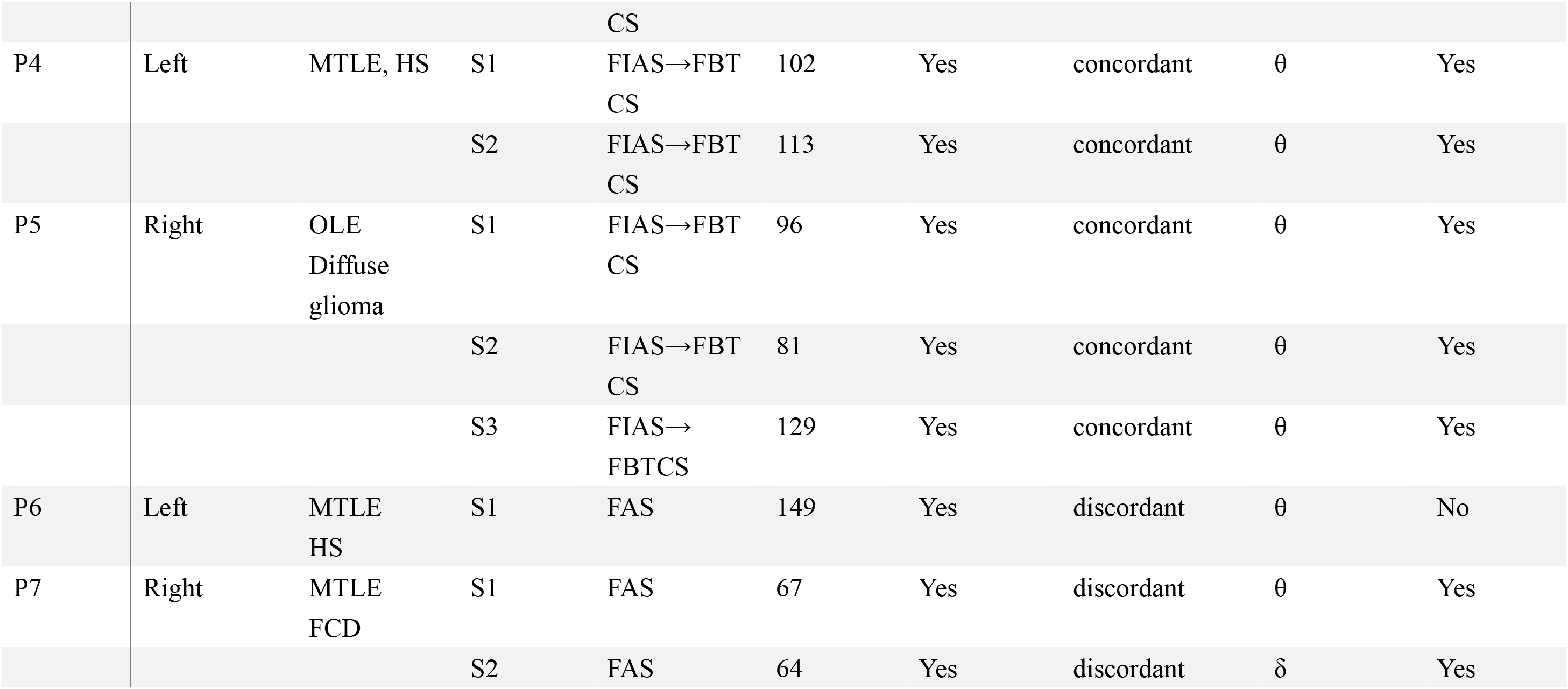
Clinical profile. SOEC: seizure onset electrode contact, PREC: preceding ripple electrode contact, MTLE: mesial temporal lobe epilepsy, DNT: dysembryoplastic neuroepithelial tumor, PLE: parietal lobe epilepsy, FCD: focal cortical dysplasia, OLE: occipital lobe epilepsy, HS: hippocampal sclerosis, FIAS: focal impaired awareness seizure, FBTCS: focal to bilateral tonic-clonic seizure, FAS: focal aware seizure *Focal resection surgery had not been performed because the detection of the seizure onset zone was impossible.

### Intracranial electrodes and their location

Data on iEEG were acquired using a combination of subdural grids (10, 20, or 30 contacts), strips (four or six contacts), and depth electrodes (six contacts) (Unique Medical Co. Ltd., Tokyo, Japan), placed using conventional craniotomy (supplemental methods). Three-dimensional (3D) brain renderings were created using FreeSurfer (https://surfer.nmr.mgh.harvard.edu) with the preoperative MRI images. Using Brainstorm (http://neuroimage.usc.edu/brainstorm/), the post-implantation CT images were overlaid onto the 3D brain renderings to obtain the position of contact for each electrode in the Montreal Neurological Institute coordinates system.

### Data acquisition and preprocessing

Signals from the iEEG were acquired at a sampling rate of 1 kHz and a time constant of 10 s, using a 128-channel digital EEG system (EEG 2000; Nihon Kohden Corporation, Tokyo, Japan), and details of preprocessing were indicated in supplemental methods. For the purpose of signal analysis, iEEG signal of each electrode contact was digitally re-referenced to a common average of all electrode contacts in each patient.

All the subsequent signal analysis was performed using MATLAB R2019b (MathWorks, Natick, MA, USA). Our iEEG data were saved every 60 min and thus each iEEG dataset contains a 60-min signal. A bandpass filter using a two-way least-squares finite impulse response filter (pop_eegfiltnew.m from the EEGLAB toolbox, https://sccn.ucsd.edu/eeglab/index.php) was applied to the preprocessed signals of the whole 60-min data to prevent edge-effect artifacts.

### High frequency activity power changes

The SO and the corresponding electrode contact (seizure onset electrode contact, SOEC) were determined by visual inspections of iEEG signals ^20^. To simplify the analysis, we randomly selected one electrode contact that fulfil the condition that it was commonly involved at seizure onset across all seizures (for patients with more than one seizure). We analyzed the iEEG data acquired 5 minutes before and after the SO for each recorded seizure. We used the ripple band (80–250 Hz) to represent HFA. The time series of the HFA power on each contact was constructed every second from the preprocessed iEEG signal by using a band-pass filter (80–250 Hz) in combination with the Hilbert transformation ^21^. The HFA power was then normalized by dividing the power at each second by the average HFA power of the initial 60 s.

### PAC analyses

We used synchronization index (SI) ^21^ to measure the strength of cross-frequency coupling between the amplitude of ripple and the phases of lower frequency bands, which include δ (1– 4 Hz), θ (4–8 Hz), α (8–13 Hz), and β (13–30 Hz).

Hilbert transformation was performed on the bandpass filtered signals to obtain the complex-valued analytic signals of each frequency band, Z_ω_(t) (ω means the frequency band). For each frequency band, the amplitude, A_ω_(t), and phase, φ_ω_(t), were calculated from the complex-valued signals using Equation 1.

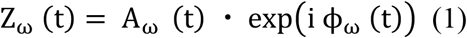

The phase of each lower frequency band, φ_l_(t), was obtained from the angle of the Hilbert transform of the bandpass filtered signal.

To obtain the surrogate signal that represent the time series of the ripple band amplitude, the amplitude of ripple band was first extracted using the squared magnitude of Z_γ_(t), the analytic signal calculated using the Hilbert transformation (P_γ_(t) = real [Z_γ_ (t)]^2^ + image [Z_γ_ (t)]^2^); then the phase of this amplitude was computed using Hilbert transformation (φ_γ_(t) = arctan (image [Z (P_γ_(t))]/real[Z(P_γ_(t))])).

SI was calculated using Equation 2.

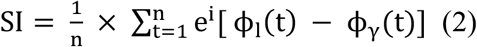

We calculated SI for every 1-s time window, which sequentially shifted every 33 ms from 5 minutes before to 5 minutes after the SO. n is the number of time points within each 1-s time window. SI is a complex number; therefore, we used the magnitude of SI, referred to as SIm. SIm varies between 0 and 1, with 0 indicating that phases are completely desynchronized and 1 indicating that phases are perfectly synchronized.

### Correlation analysis related to PAC

We analyzed the correlation between SIm and ripple normalized power in the following three states in relation to seizure onset: pre-ictal (from −1.5 to 0 min before the SO), ictal (from 0 to 1.5 min after the SO), and late-ictal (from 1.5 to 3.0 min after the SO) states using implanted all contacts (total 1189 electrode contacts). Pearson correlation coefficients were calculated.

### Phase-conditioned analysis

To identify the lower frequency phase to which ripple power was coupled, we computed the average oscillation of each lower frequency band on the SOEC across all seizures (15 seizures) and the average ripple normalized power on the SOEC. The phases of δ and θ band, were divided into 12 intervals of 30° without overlaps: 0° ± 15°, 30° ± 15°,…, 300° ± 15°, and 330° ± 15°, resulting in 12 phase bins. The reason for choosing these two bands is described in the result section. For each state (pre-ictal, ictal, and late-ictal), the ripple normalized power on the SOEC was averaged within each phase bin. Using signal from the SOEC instead of the average of all electrode contacts avoids averaging from obscuring the characteristics-of-interest.

### Visualization of synchronized multimodal data (Video)

Multimodal data, including iEEG waveform, ripple normalized power, and SIm, were synchronized and simultaneously displayed. To construct a power distribution map, we plotted red circles on the brain image to indicate the electrode locations, and the diameters were scaled linearly with ripple power (supplemental methods).

### Statistics

The ripple normalized power and SIm values were averaged across all electrode contacts in each seizure, and then averaged ripple normalized power and averaged SIm values were averaged across all seizures (Fig. 1A, 1B, and 1C). SIm values were normalized using SIm values calculated from the data acquired five minutes before SO to allow comparison between different low frequency bands (Fig. 1B). In order to know if there is a difference between frequency band with the highest SIm and the others, we used single-sided Wilcoxon signed-rank test for pairwise comparison between frequency bands and Bonferroni correction for multiple comparisons (Fig. 1A, 1B, and 1D). To define the change of the power and SIm (Fig. 1C) compared to the background (the initial 10-s data), we used a permutation test for the comparison between the initial 10-s data and the next sequential 10-s data ^22^ and a familywise error (FWE)-corrected threshold for multiple comparisons (supplemental methods). The values above the FWE-corrected threshold are statistically significant ^23^.

**Figure 1.**
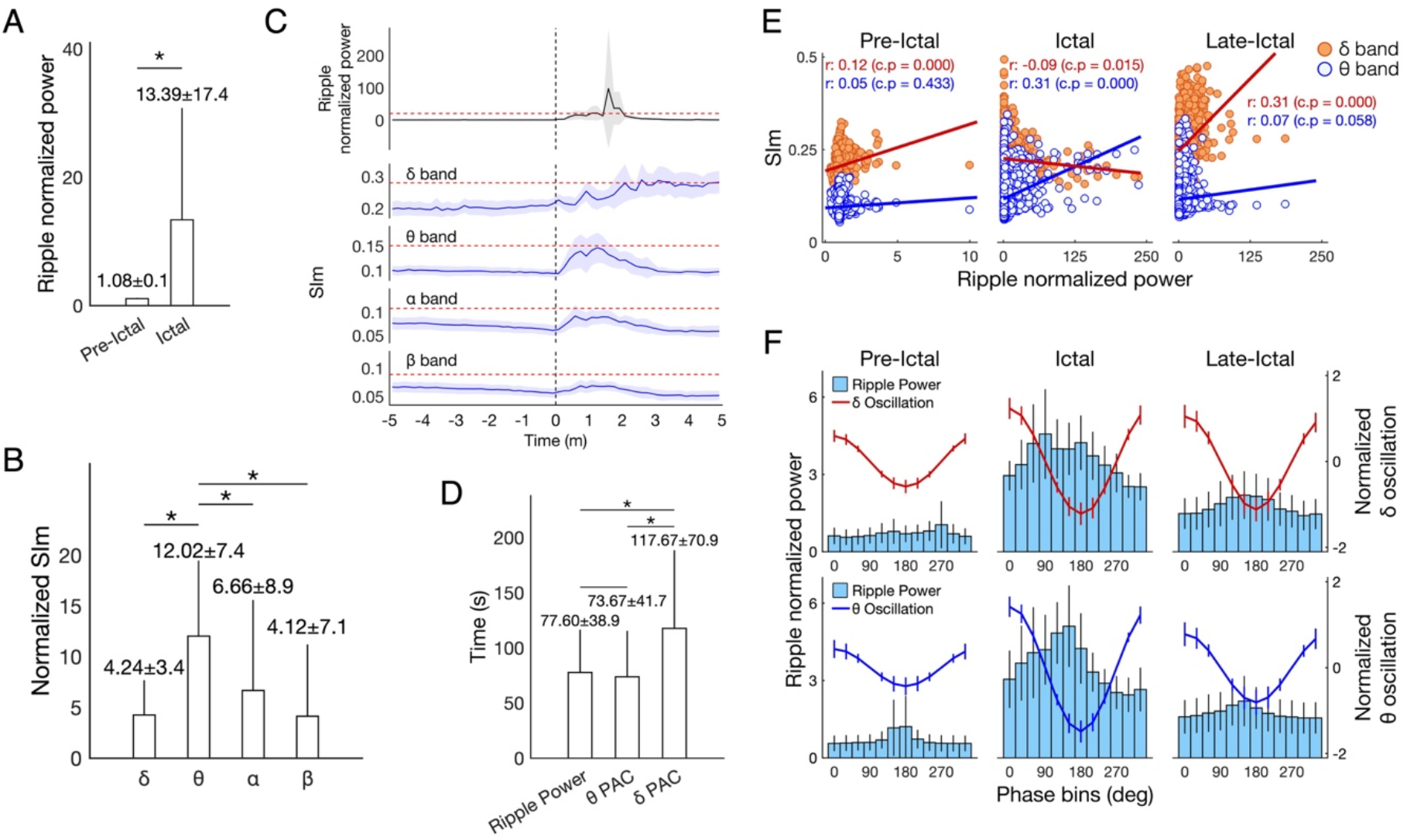
Characteristics of ripple power and PAC related to seizures. A. The ripple normalized power in ictal state was significantly greater than that in pre-ictal state. B. The normalized SIm during ripple power burst related to seizures achieved the significant highest values in θ band. C. Temporal plot of ripple normalized power, θ-, α-, and β-SIm increased and achieved the peak after the SO (0 min). δ-SIm achieved the peak after completion of ripple power burst. The FWE-corrected threshold is indicated as red dashed line. D. The time taken to achieve the maximum values was compared between ripple normalized power, θ-SIm (PAC), and δ-SIm (PAC). The time of δ PAC was significantly slower. E. During ictal state, significant positive correlation between ripple normalized power and θ-SIm, and significant negative correlation between ripple normalized power and δ-SIm were observed. In late-ictal state, significant positive correlation between ripple normalized power and δ-SIm was observed. r: correlation coefficients, c.p: corrected p value with Bonferroni correction. F. Phase tuning δ and θ oscillation showed the trough at 180°. The phase-tuning ripple normalized power showed the peak at the trough with θ phase in ictal state, and with δ phase in late-ictal state. The error bars (EB) in 1A, 1B, and 1D indicate standard deviation. The EB in 1C, and 1F indicate 95% confidence intervals.

To assess the significant change in SIm, we used the boot-strapping technique (Fig. 2–4, and Video S1‒S3) ^21^ (supplemental methods). To correct for multiple comparisons, we used the FWE-corrected threshold (95%) ^23^.

**Figure 2.**
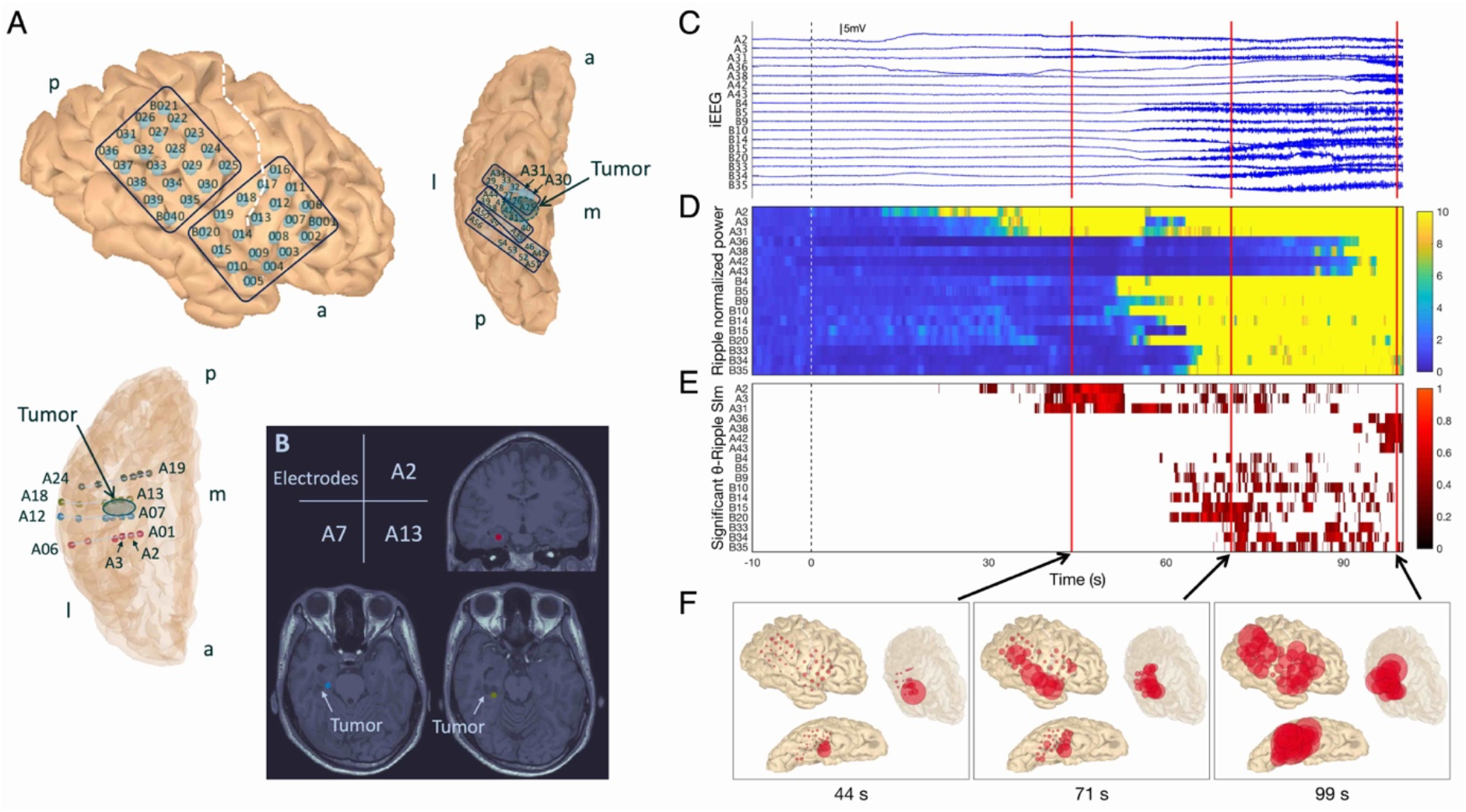
Seizure profile in Patient 1 (P1)-Seizure 1 (S1). A. Depth electrodes A1–6, A7–12, A13–18, and A19–24 were targeting the parahippocampal gyrus, the anterior region of the tumor, the posterior region of the tumor, and the lingual gyrus, respectively. a. anterior; p. posterior; m. medial; l. lateral. B. The location of A2, A7, and A13 in relation the cystic tumor is shown on the T1-weighted MRI. C. Intracranial electrodes’ raw signals. Initial infraslow activities were observed on A2. D. The ripple normalized power started to increase after the SO, from the A2 and propagated to other contacts. E. Significant θ-ripple SIm shows the cluster in contacts in which ripple power increased. Here, 0 s corresponds to the SO determined visually and is indicated as dashed lines. F. Ripple power distribution map. The location of red circles corresponds to each contact’s location, and those located within the semitransparent brain were depth electrode contacts. The size of red circles corresponds to ripple power. The ripple power started to increase in A2 (44 s) and propagated to other contacts on the cortex along the Sylvian fissure (71 s), and almost all other regions (99 s).

## Results

### Profile of ripple power and PAC changes related to seizures

We compared ripple normalized power between the pre-ictal and the ictal states. The ripple normalized power during seizures (ictal state) were significantly larger than that before seizures (pre-ictal state) (single-sided Wilcoxon signed-rank, P = 3.1 × 10^−5^) (Fig. 1A). Averaged time of seizures is 125.73 ± 55.1 s, and times of each seizure are shown in Table 1.

Among all seizures analyzed, SOEC that was determined by conventional visual inspection were concordant with the preceding ripple electrode contact (PREC, see case studies below) that was determined by ripple activities in 12/15 seizures (80%). When focusing on only focal to bilateral tonic-clonic seizures (FBTCS), SOEC matched PREC in all seizures (Table 1).

We compared SIm values calculated from the coupling between ripple and each lower frequency band during the burst of ripple power within the seizure state. The normalized SIm of ripple-θ was the highest among all (single-sided Wilcoxon signed-rank with Bonferroni correction, θ-δ: corrected P = 3.0 × 10^−3^, θ-α: corrected P = 1.3 × 10^−2^, θ-β: corrected P = 2.3 × 10^−3^) (Fig. 1B). The main low frequency band coupled with ripple was the δ band in 2/15 seizures (13%), θ band in 12/15 seizures (80%), and α band in 1/15 seizures (7%) (Table 1).

### Temporal profile of ripple power and PAC changes

Dynamic changes of ripple normalized power and PAC between ripple and each lower frequency band from 5 min before to 5 min after SO are shown in Fig. 1C. A significant burst of ripple power was observed after SO, which was accompanied by two different profiles of PAC with the lower frequency bands: (1) PAC changes with a peak (θ-, α-, and β-ripple coupling) and (2) PAC changes with a gradual increase and a plateau (δ-ripple coupling). For those with the first profile, the PAC for θ band after SO were the higher than those for α and β bands. For the second profile, the PAC for δ band increased gradually and reached its maximum after ripple power burst. We focused on PAC for θ band (representing the first profile) and δ band (the second profile) in the following analyses.

There was no significant difference between the time taken (since SO) to attain maximum ripple power (77.6 ± 38.9 s) and maximum θ PAC (73.7 ± 41.7 s); however, the time taken to achieve maximum value of δ PAC (117.7 ± 70.9 s) was significantly more than that for both ripple power and θ PAC (single-sided Wilcoxon signed-rank with Bonferroni correction, ripple power and δ PAC, corrected P = 1.6 × 10^−3^, θ PAC and δ PAC, corrected P = 8.2 × 10^−4^) (Fig. 1D). A time lag between θ and δ PAC was observed in 14/15 seizures (93.3%) (Table 1).

### Correlation between ripple normalized power and SIm

To clarify the differences between PAC for θ and δ bands, correlation analysis was used. The correlation of ripple with the SIm of θ band was positive in all three states (pre-ictal, ictal and late-ictal), in which a statistical significance was reached in the ictal state. In contrast, the correlation with the SIm of δ band was positive in the pre- and late-ictal states and negative in the ictal state, in which a statistical significance was reached in all three states (Fig. 1E).

### Phase-tuning ripple power

The oscillation of δ and θ bands showed a trough at 180° in all pre-ictal, ictal and late-ictal states (Fig. 1F). In the ictal state, in which θ-SIm increased (Fig. 1C), ripple normalized power peaked at the trough of the θ oscillation but not the δ oscillation (Fig. 1F). In the late-ictal state, in which δ-SIm increased (Fig. 1C), ripple normalized power peaked at the trough of the δ oscillation (Fig. 1F).

### Case studies

During seizures, the ripple power increased and was coupled with the θ phase. After completion of ripple power burst, δ-ripple coupling occurred. We show the synchronized multimodal data including iEEG waveforms, ripple normalized power, SIm, and ripple power distribution map in the figure and videos of three illustrative cases. In Fig. 2, 3, and 4, significant θ-ripple SIm and representative electrodes are shown. In supplementary videos (Video S1–3), significant δ- and θ-ripple SIm and all electrodes are shown.

**Figure 3.**
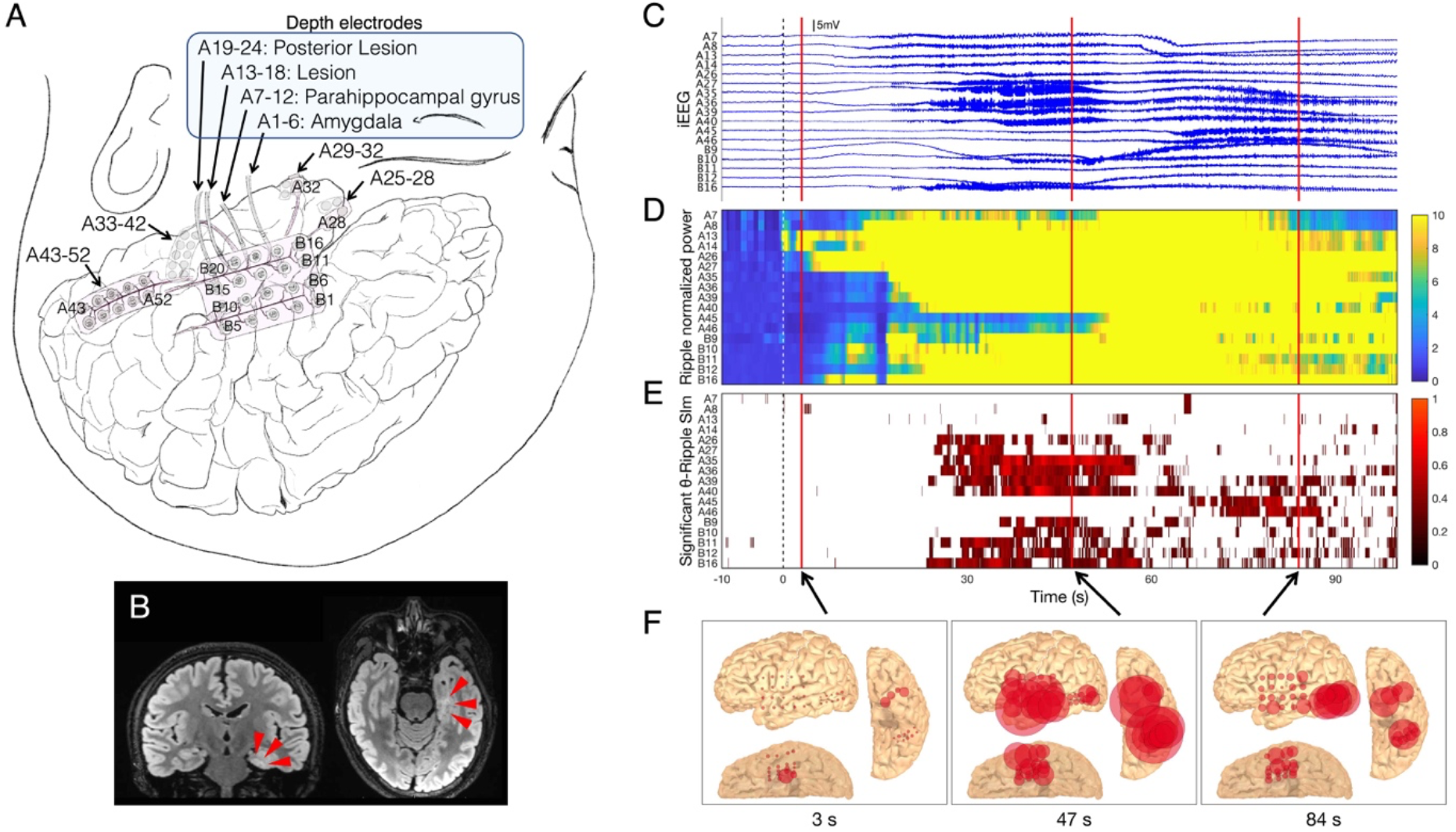
Seizure profile in P4-S2. A. Depth electrodes A1–24 were inserted into the left mesial temporal lobe. A13 was located in the lesion. B. The high-intensity lesion in the left mesial temporal lobe is shown on FLAIR MRI (red wedge arrows). C. In A13, initial infraslow activities and low-voltage fast waves were observed. D. The ripple normalized power started to increase after the SO, from A13 and propagated to other contacts. E. Significant θ-ripple SIm shows the cluster in contacts in which ripple power increased. Here, 0 s corresponds to the SO determined visually and is indicated as dashed lines. F. Ripple power distribution map. The location of red circles corresponds to each contact’s location, and those located within the semitransparent brain were depth electrode contacts. The size of red circles corresponds to ripple power. The ripple power started to increase in A13 (3 s) and propagated to other contacts on the cortex along the Sylvian fissure (47 s), and to the posterior middle temporal gyrus (84 s).

**Figure 4.**
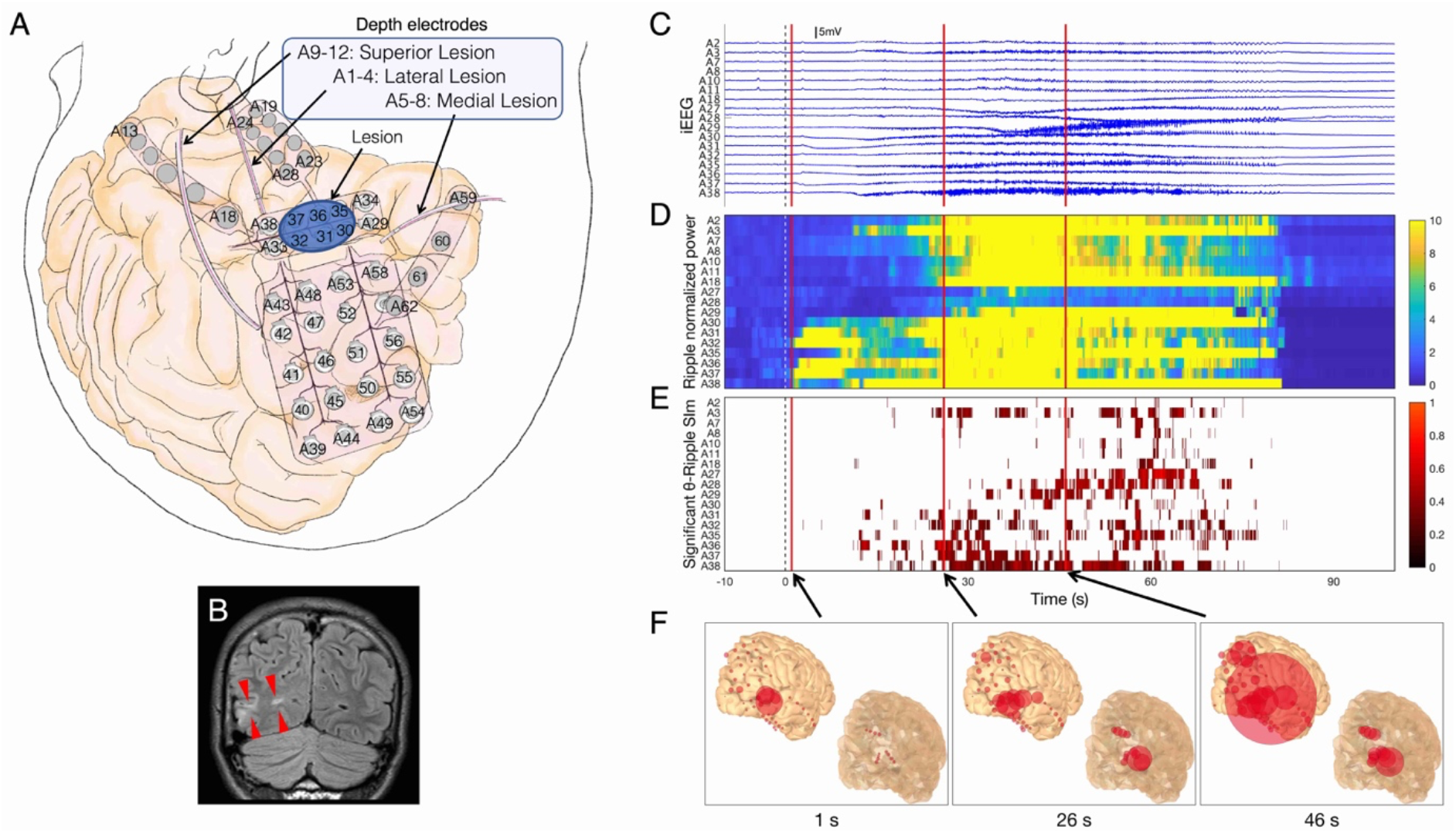
Seizure profile in P5-S2. A. Depth electrodes A1–12 were targeting around the right occipital lesion. B. The high-intensity lesion in the right occipital lobe is shown on FLAIR MRI (red wedge arrows). C. In A32 and A37, initial infraslow activities and low-voltage fast waves were observed. D. The ripple normalized power started to increase after the SO, from the A32 and A37 and propagated to other electrodes. E. Significant θ-ripple SIm shows the cluster in contacts in which ripple power increased. Here, 0 s corresponds to the SO determined visually and is indicated as dashed lines. F. Ripple power distribution map. The location of red circles corresponds to each contact’s location, and those located within the semitransparent brain were depth electrode contacts. The size of red circles corresponds to ripple power. The ripple power started to increase in A32 and A37 over the right occipital lesion (1 s) and propagated to the parietal lobe (46 s).

### Case #1 (P1-S1 in Table 1)

Interictal scalp EEG recorded spike-wave complexes over the right temporal region. We placed intracranial electrodes in the right hemisphere (Fig. 2A). MRI images showed a cystic lesion in the right mesial temporal lobe (MTL) (Fig. 2B).

We captured electroclinical seizures consisting of focal impaired awareness seizure (FIAS), followed by FBTCS (Fig. 2 and Video S1). Contact A2 on the depth electrode inserted in the right parahippocampal gyrus, showed initial infraslow activities, followed by low-voltage fast waves that changed into high-amplitude fast waves and spread to the other electrodes (Fig. 2C). Ripple power increases also began from A2 and spread to the other electrodes (Fig. 2D and 2F), and in this case, the SOEC was concordant with the PREC. Significant θ-ripple SIm values were observed in the electrodes in which ripple power increased (Fig. 2E), and after that, significant δ-ripple SIm values were observed (Video S1). The time lag between θ and δ PAC was observed. Ripple-band power (represented by the size of the red circle) fluctuated rhythmically at certain rhythms when θ-SIm reached statistical significance and this relationship was also observed for δ-band (Video S1).

### Case #2 (P4-S2 in Table 1)

Interictal scalp EEG recorded spike-wave complexes over the left frontotemporal region. We placed intracranial electrodes in the left hemisphere (Fig. 3A). MRI fluid-attenuated inversion recovery (FLAIR) images showed a high-intensity lesion in the left MTL (Fig. 3B).

We captured electroclinical seizures consisting (Fig. 3 and Video S2). Contact A13 on the depth electrodes, which were inserted in the left mesial temporal lesion, showed initial infraslow activities, followed by low-voltage fast waves that changed into high-amplitude fast waves and spread to the other electrodes (Fig 3C). Ripple power increases also began from A13 and propagated to other electrodes (Fig. 3D and 3F), and in this case, the SOEC was concordant with the PREC. Significant θ-ripple SIm values were observed in contacts in which ripple power increased (Fig. 3E), and after that, significant δ-ripple SIm values were observed (Video S2). A time lag between θ and δ PAC was observed. Rhythmic ripple-band activities were visualized in the power distribution map when the values of SIm were significantly high (Video S2).

### Case #3 (P5-S2 in Table 1)

Interictal scalp EEG recorded spike-wave complexes over the right temporo-occipital region. We placed intracranial electrodes in the right hemisphere (Fig. 4A). MRI FLAIR images showed a high-intensity lesion in the right occipital lobe (Fig. 4B).

We captured electroclinical seizures, followed by bilateral lower extremities stereotypies (FBTCS) (Fig. 4 and Video S3). The A32 and A37 electrodes, which were placed over the occipital lesion, showed initial infraslow activities, followed by low-voltage fast waves that changed into high-amplitude fast waves (Fig 4C). Ripple power increases also began from A32 and A37 electrodes and propagated to the other electrodes (Fig. 4D and 4F), and the SOEC was concordant with the PREC. Significant θ-ripple SIm values started to appear in electrodes which were placed over the occipital lesion (Fig. 4E), and after that, significant δ-ripple SIm values were also observed (Video S3). The time lag between θ and δ PAC was observed. Rhythmic ripple-band activities were visualized in the power distribution map when the values of SIm were significantly high (Video S3).

## Discussion

This study demonstrated that a ripple power burst occurs during seizures, and the phase of θ band modulates ripple power involved in seizures. Our videos showed that when ripple power increased, the individual ripple power of each electrode contact (corresponded to the size of each red circle in the video) changes rhythmically. During such rhythmic fluctuation in the circles’ size, a significant θ-ripple PAC was observed. Therefore, we inferred that the rhythmic fluctuation in the circles’ size (in the video) was modulated by the θ rhythm and this fluctuation represented the θ-ripple PAC phenomenon. Our video is a useful tool for understanding visually and comprehensively the PAC phenomenon related to ripple activities during seizures.

How HFAs were propagated during seizures has been visualized by topographic videos ^6^ ^24^, however, PAC changes involved in seizures have not been visualized. In our videos, the diameters of each circle that represents an implanted electrode contact changed with ripple power, and the videos also demonstrated the propagation of HFAs by dynamic changes of circles’ diameters. Moreover, the rhythmic fluctuations of the diameter were observed in each contact. When δ- or θ-ripple PAC significantly increased, the rhythmic fluctuation of the circles’ diameter became especially obvious. We inferred that this rhythmic fluctuation of the circles’ diameter was tuned at δ- or θ rhythm, and concluded that this rhythmic fluctuation enabled the visualization of the PAC phenomenon.

HFAs have been observed during seizures ^24^ ^25^ and HFOs are suggested as useful biomarkers for detection of the SOZ ^26^ ^27^. In this study, significant increase in ripple power was observed during seizures, and PREC which showed preceding ripple activities were demonstrated (Fig. 2D, 3D, and 4D, and supplementary videos). The SOEC was conventionally determined by visual inspection ^20^, and we demonstrated that in FBTCS, the SOECs were concordant with the PRECs. Therefore, we thought that ripple activities were useful for detection of the SOZ.

In line with a previous study ^14^, our study demonstrated that θ band was the main low frequency band modulating ictal-ripple activities and α- and β-ripple PAC showed the same tendency as the θ-ripple PAC with a weaker PAC though. Moreover, it is reported that coupling with θ waves and HFOs well discriminated normal brain regions from SOZ ^28^. Therefore, we inferred that ripple activities occurring during early seizure evolution were modulated by θ rhythm. The occurrence of maximal ripple power at the trough of the low frequency oscillation was concordant with a previous study^14^. In an interictal state, a positive correlation between coupling and ripple amplitude were known^29^, and we showed the positive correlation between θ-ripple PAC and ictal-ripple power during the ictal state.

In contrary, we showed newly finding that δ-ripple PAC had a negative correlation with ripple power during ictal-ripple power burst, and increased after ictal-ripple power burst subsided. Coupling between HFAs and δ band were investigated in previous studies associated with epileptic spasm^18^ and an interictal state^30^. Our videos brought a new insight that δ-ripple PAC significantly increased after θ-ripple PAC. Our results suggested that δ- and θ-ripple PAC were caused by different mechanisms, and this explained the time lag between δ- and θ-ripple PAC.

This study had some limitations. Because intracranial EEG electrodes covers only a small portion of the brain, we can never be sure if the intracranial electrodes were placed in the actual SOZ, and thus the activities that we analyzed may not be the actual activities from the SOZ. To limit the effect of this uncertainty, we used the average ripple power and SIm from all electrodes because we could evaluate at least seizure-related changes which were propagated activities from the SOZ not focal activities of the SOZ. All the seizures were recorded after an extensive reduction of antiepileptic drugs, and thus, they might not represent the patients’ usual seizures. However, our analysis were independent from this issue because reduction in medication does not affect the morphology of discharges at onset, and duration of contralateral spread ^31^. Because this study included only patients with focal epilepsy, our findings may not be generalized to patients with generalized epilepsy.

## Acknowledgements

This study was supported by the Grants-in-Aid for Scientific Research (A) (KAKENHI; grant no., 18H04166) and the Grants-in-Aid for Early-Career Scientists (KAKENHI; grant no., 18K18366), which are funded by the Japan Society for the Promotion of Science (JSPS; Tokyo, Japan).

## Author Contributions

H.H. conceived the study, collected the data, created the MATLAB program, analyzed the data, created all figures and the video, and was primarily responsible for writing the manuscript. H.M.K., N.T., S.O., H.K., and M.H. performed the epileptic surgery. All authors clinically cared for and evaluated the patient. H.M.K., T.Y., and M.H. advised H.H. on scientific matters. H.M.K revised the manuscript. H.K. and M.H. supervised this study. All authors have reviewed the manuscript.

## Conflict of Interests

None of the authors has any conflict of interest to disclose.

**Common Legend for Video**

Multimodal data are shown simultaneously, and each file includes ripple (80–250 Hz) power distribution map, iEEG signals, ripple normalized power, θ-ripple PAC (SIm), and δ-ripple PAC (SIm). The video starts 10 s before the SO. In power distribution map, red circles, corresponding to intracranial electrode contacts, were scaled linearly with ripple power changes. The circles in the semitransparent brain indicate depth electrode contacts. The vertical bars indicating current-time are colored red in iEEG signals, white in ripple normalized power, and blue in PAC (SIm). SIm values scaled as black to red are statistically significant values to which the FWE-corrected threshold was applied. All implanted electrode contacts after removal of noisy contacts are shown in the vertical axes.

**Legend for Video S1**

The seizure in P1-S1 is shown. The rhythmic movement of red circles started from the A2 depth electrode contact, which were inserted into the right parahippocampal gyrus. The rhythmic fluctuations spread to the contacts over the cortex along the Sylvian fissure and that over the temporal base. During significantly high values of SIm, the fluctuation of red circles changed at certain rhythms. We inferred that the rhythmic fluctuations were tuned at θ or δ rhythm, and represented the PAC phenomenon. Time lag between θ and δ SIm were observed. The SO is 11:22:03. The 44 s, 71 s, and 99 s in Fig. 2 correspond to 11:22:47, 11:23:14, and 11:23:42 in respectively.

**Legend for Video S2**

The seizure P4-S2 were shown. The rhythmic movement of red circles started from the A13 depth electrode contact, which were inserted into the left mesial temporal lesion. The rhythmic movement spread to the contacts over the temporal tip cortex and that over the cortex along the Sylvian fissure and that over the posterior middle temporal gyrus. Time lag between θ and δ SIm were observed. The SO is 4:15:56. The 3 s, 47 s, and 84 s in Fig. 3 correspond to 4:15:59, 4:16:43, and 4:17:20 in respectively.

**Legend for Video S3**

The seizure P5-S2 were shown. The rhythmic movement of red circles started from the A32 and 37 surface electrode contacts placed over the right occipital lesion. The rhythmic movement spread to the contacts over the parietal lobe. Time lag between θ and δ SIm were not observed. The SO is 8:28:53. The 1 s, 26 s, and 46 s in Fig. 4 correspond to 8:28:54, 8:29:19, and 8:29:39 in respectively.

## Supplemental material

## Methods

### Intracranial electrodes and their location

The diameter of each contact was 3 or 5 mm, and the inter-contact distance was 5, 7, or 10 mm for grid and strip electrodes. The diameter was 1.5 mm, and the inter-contact distance was 5 mm for depth electrodes.

### Data preprocessing

The raw signals of iEEG were then preprocessed using a low-pass filter at 333 Hz (to prevent aliasing) and a 60-Hz notch filter (to eliminate the AC line artifact) using the BESA Research 6.0 software (BESA GmbH, Grafelfing, Germany). Artifactual signals from electrodes were excluded from further analyses.

### Visualization of synchronized multimodal data (Video)

To construct a power distribution map, we calculated the power of ripple using the periodogram. The period of 10 seconds before SO was defined as the baseline, and the baseline ripple power was obtained from the baseline segment. We calculated the ripple power within a 500-ms sequential time window every 33 ms (30 fps). The ripple power ratio was calculated by dividing the ripple power of the 500-ms time window with baseline ripple power. We plotted red circles on the brain image to indicate the electrode locations, and the diameters were scaled linearly with ripple power ratio.

### Statistics

To define the change of the power and SIm (Fig. 1C) compared to the background (the initial 10-s data), we used a permutation test for the comparison between the initial 10-s data and the next sequential 10-s data, and a familywise error (FWE)-corrected threshold for multiple comparisons. Each permutation test produces a set of differences between the initial 10-s data and the next sequential 10-s data. The maximum value of the differences from each permutation test were stored. The values at 95% of the distribution of these maximum values were taken as the FWE-corrected threshold.

To assess the significant change in SIm, we used the boot-strapping technique (Fig. 2–4, and Video S1‒S3). First, the phase of ripple power time series was shifted in time by a random amount. Then, this phase-shifted ripple power time series was used to calculate a SIm value for the purpose of bootstrapping (SImb). For each pair of ripple power and a lower frequency band amplitude, this procedure was repeated 1000 times to create the distribution of SImb^1^.

## Notes

### Competing Interest Statement

The authors have declared no competing interest.

### Summary of Updates

The figures were revised for anonymity.

